# Efficacy of Host Cell Serine Protease Inhibitor MM3122 against SARS-CoV-2 for Treatment and Prevention of COVID-19

**DOI:** 10.1101/2024.02.09.579701

**Authors:** Adrianus C.M. Boon, Traci L. Bricker, Ethan J. Fritch, Sarah R. Leist, Kendra Gully, Ralph S. Baric, Rachel L. Graham, Brigid V. Troan, Matthew Mahoney, James W. Janetka

## Abstract

We have developed a novel class of peptidomimetic inhibitors targeting several host cell human serine proteases including transmembrane protease serine 2 (TMPRSS2), matriptase and hepsin. TMPRSS2 is a membrane associated protease which is highly expressed in the upper and lower respiratory tract and is utilized by SARS-CoV-2 and other viruses to proteolytically process their glycoproteins, enabling host cell receptor binding, entry, replication, and dissemination of new virion particles. We have previously shown that compound MM3122 exhibited sub nanomolar potency against all three proteases and displayed potent antiviral effects against SARS-CoV-2 in a cell-viability assay. Herein, we demonstrate that MM3122 potently inhibits viral replication in human lung epithelial cells and is also effective against the EG.5.1 variant of SARS-CoV-2. Further, we have evaluated MM3122 in a mouse model of COVID-19 and have demonstrated that MM3122 administered intraperitoneally (IP) before (prophylactic) or after (therapeutic) SARS-CoV-2 infection had significant protective effects against weight loss and lung congestion, and reduced pathology. Amelioration of COVID-19 disease was associated with a reduction in pro-inflammatory cytokines and chemokines production after SARS-CoV-2 infection. Prophylactic, but not therapeutic, administration of MM3122 also reduced virus titers in the lungs of SARS-CoV-2 infected mice. Therefore, MM3122 is a promising lead candidate small molecule drug for the treatment and prevention of infections caused by SARS-CoV-2 and other coronaviruses.

**IMPORTANCE:** SARS-CoV-2 and other emerging RNA coronaviruses are a present and future threat in causing widespread endemic and pandemic infection and disease. In this paper, we have shown that the novel host-cell protease inhibitor, MM3122, blocks SARS-CoV-2 viral replication and is efficacious as both a prophylactic and therapeutic drug for the treatment of COVID-19 in mice. Targeting host proteins and pathways in antiviral therapy is an underexplored area of research but this approach promises to avoid drug resistance by the virus, which is common in current antiviral treatments.

## INTRODUCTION

The COVID-19 pandemic, caused by the Severe Acute Respiratory Syndrome Coronavirus 2 (SARS-CoV-2), has illuminated the devastating impact of new emerging viral diseases on both the global economy and health of populations^1^. It has affected all aspects of human life and highlighted the threat of future outbreaks with other respiratory viruses for which currently available drugs will be ineffective. Incredibly, vaccines as well as several new drug candidates targeting this virus were developed at unprecedented speed because of early countermeasure work that focused on paradigm pathogens within the coronavirus family^2–4^. This was made possible by the combined efforts of scientists worldwide who elucidated details about the makeup and pathogenesis of SARS-CoV-2 infection. This work also resulted in the identification of several potential therapeutic targets such as the viral entry receptor, angiotensin converting enzyme 2 (ACE2), and host cell proteases, such as transmembrane protease serine 2 (TMPRSS2) and cathepsin L1 (CTSL1), required for entry, replication, and release of the virus^5–8^.

TMPRSS2^9–11^ has a trypsin-like serine protease domain and belongs to the family of Type II Transmembrane Serine Protease (TTSP) proteolytic enzymes with reported physiological roles in cancer and many other diseases^12–15^. TMPRSS2 has previously been shown to be important in other coronavirus infections caused by SARS-CoV-1^16, 17^, HKU-1^18^, MERS-CoV^19^, and others^20,21^. Furthermore, both TMPRSS2 and the other TTSPs, matriptase (ST14)^22, 23^ and HAT (human airway trypsin-like protease, TMPRSS11D)^24, 25^, have been demonstrated to proteolytically process the hemagglutinin (HA) protein on the surface of some influenza A viruses and SARS-CoV-1^17^, allowing viral cell adhesion and entry in these infections^24, 26–31^. Additionally, TMPRSS2 was found to support replication of other respiratory viruses including human para-influenza virus type 1, 2 and mouse Sendai virus^32^. This makes TMPRSS2 an excellent target for the development of a broadly acting antiviral inhibitor^33^ against diverse respiratory viruses^24, 34–39^.

We recently reported on the discovery and development of a new class of peptidomimetic TMPRSS2 inhibitors^39^. These inhibitors were rationally designed based on the peptide substrate sequence specificity of TMPRSS2^15^ and molecular docking studies using the X-ray structure of TMPRSS2 bound to another inhibitor nafamostat^40, 41^. These inhibitors, including MM3122 and MM3144, are significantly more potent than another reported TMPRSS2 inhibitor Camostat^41^, suggesting improved efficacy *in vivo*. Unlike MM3122, which has the desired selectivity for TMPRSS2 over the coagulation serine proteases thrombin and Factor Xa, MM3144 does not. This compound was subsequently reported as a matriptase and TMPRSS2 inhibitor by another group as N-0386^42^. We tested the *in vitro* and *in vivo* efficacy of the most promising compound at the time, MM3122, in Calu-3 human lung epithelial cells and in a mouse model of SARS-CoV-2. Overall, we showed potent antiviral efficacy of MM3122 against XBB.1.5 and EG.5.1 variant of SARS-CoV-2 *in vitro* and amelioration of COVID-19 disease *in vivo*.

## RESULTS

### MM3122 inhibits authentic SARS-CoV-2 replication in human lung epithelial cells

The ability of MM3122, a TMPRSS2 inhibitor, to inhibit wild-type (wt) SARS-CoV-2 infection and replication was assessed on Calu-3 cells, a human lung epithelial cell line. At a 0.03 µM concentration of MM3122, virus replication was completely inhibited, and no infectious virus was detected in the supernatant of the treated and SARS-CoV-2 infected cells (**Fig 1**). The inhibitory concentration (IC_50_) of MM3122 was ∼0.01-0.02 µM against the authentic wt SARS-CoV-2 virus. This is greater than 50 times more potent than Remdesivir which had an IC_50_ of ∼1 µM, an RNA-dependent RNA polymerase inhibitor developed for other viral infections that is one of the FDA-approved drugs approved to treat SARS-CoV-2 infected patients^43^. MM3122 was also tested for its activity against the EG.5.1 variant of SARS-CoV-2. Similar to wt SARS-CoV-2, we observed robust inhibition of the virus with an IC_50_ of ∼0.05-0.1 µM. Taken together, these studies demonstrate the high potential for small molecule TMPRSS2 inhibitors to inhibit replication of SARS-CoV-2 and several of its many variants.

**Figure 1:**
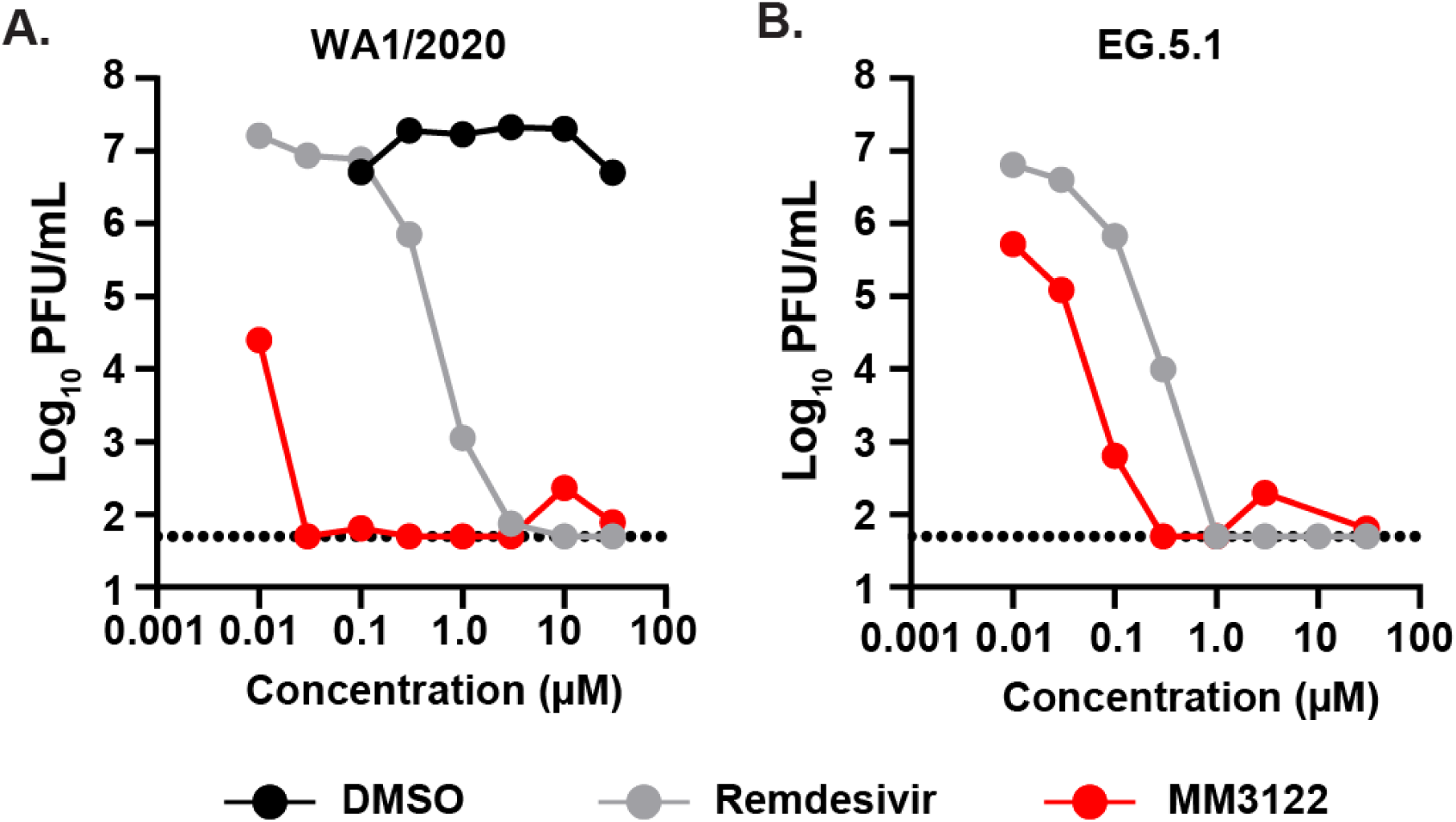
The TMPRSS2 inhibitor MM3122 inhibits replication of authentic SARS-CoV-2 and variants. Calu-3 cells, plated in 24-well plates, were infected with 4,000 PFU of A) WA1/2020 and B) EG.5.1 strains of SARS-CoV-2 for one hour at 37°C. After two washes, the cells were incubated for 48 hours in media with different concentrations of MM3122 (red symbol), Remdesivir (grey symbols), or DMSO (black symbols) as the positive and negative controls respectively. Infectious virus titer in the supernatant of the wells 48 hours after infection with SARS-CoV-2. The results are the average of 3 independently repeated assays. The dotted line is the limit of detection.

### MM3122 is a multi-targeted serine protease inhibitor with activity against some cathepsins

To determine the target specificity of MM3122 for TMPRSS2, we tested for its inhibitory activity against 53 serine and cysteine proteases (**Table 1**). In our previous paper^39^, we profiled MM3122 for its inhibition of hepatocyte growth factor activator (HGFA), matriptase, hepsin, thrombin, and Factor Xa. For the 47 other proteases we contracted Reaction Biology Co. (Malvern, PA) to determine the IC_50_ values of MM3122 against a large panel of serine and cysteine proteases of high importance. A previous group had reported the selectivity profile for Camostat and Nafamostat against these same proteases^44^ which is also shown in **Table 1**. In addition to TMPRSS2, MM3122 has potent activity (0.01 to 10 nM) against only 7 other proteases, matriptase, hepsin, matriptase-2, plasma kallikrein, trypsin, tryptase b2 and tryptase g1. It has moderate inhibitory activity (10 nM to 1 µM) against HGFA, Factor Xa, kallikrein 1 (KLK1), KLK5, KLK14, plasmin, and proteinase K and surprisingly against the cysteine protease cathepsin S with an IC_50_ of 590 nM. Furthermore, MM3122 also inhibited the other cysteine proteases cathepsin C, cathepsin L and papain, with IC_50_s of 1.4 µM, 12.8 µM, and 1.1 µM. Comparing the selectivity profile of MM3122 to that of Camostat and Nafamostat reveals that the cysteine protease activity is absent in the latter. Otherwise, the profiles are generally similar with some exceptions, notably decreased activity of MM3122 against thrombin, plasmin, Factor VIIA and XIA, KLK12, KLK13 and KLK14, urokinase, matriptase-2, trypsin and the 2 tryptases. We also found that MM3122 does not inhibit furin or the SARS-CoV-2 proteases, M_pro_ and PL_pro_.

**Table 1:**
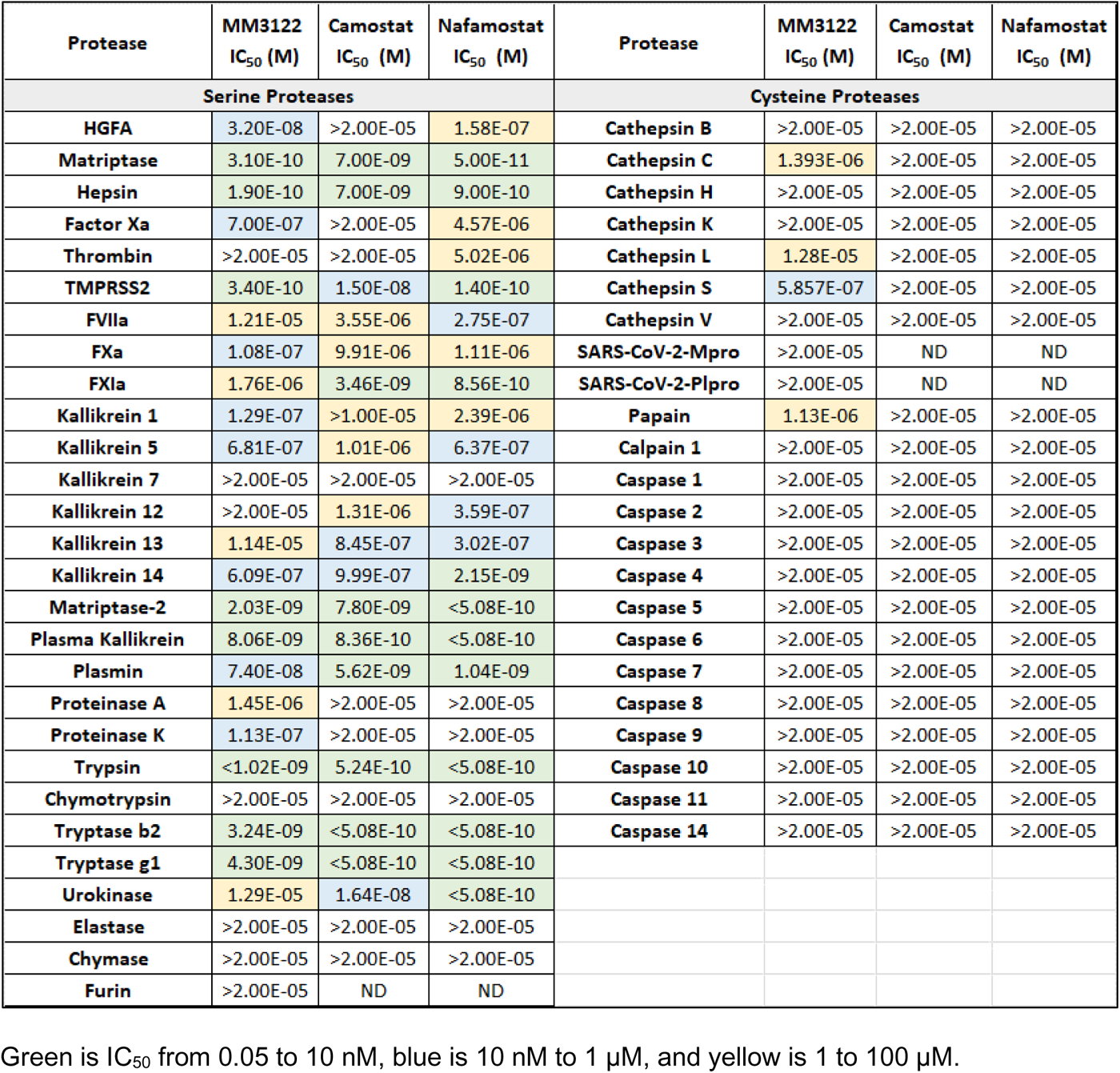
Protease selectivity profile of MM3122, Camostat and Nafamostat tested against a panel of 53 serine and cysteine proteases.

### MM3122 ameliorates SARS-CoV-2 induced disease in mice

To assess the antiviral activity of MM3122 *in vivo*, four groups of mice were treated with 50 mg/kg or 100 mg/kg MM3122 30 minutes prior to (prophylactic) and 24 hours after (therapeutic) intranasal infection with the MA10 strain of SARS-CoV-2. A mock infected and vehicle treated group were included as controls. Intranasal inoculation of vehicle treated mice with SARS-CoV-2 resulted in significant weight loss compared to mock infected animals (**Fig 2A**). Importantly, none of the MM3122 groups lost any significant amount of body weight. Five days after infection, lung congestion was assessed using an independently developed scoring system as described in the Methods. Vehicle-treated mice that were infected with SARS-CoV-2 MA10 demonstrated evidence of congestion with scores ranging from 1-3. These scores were reduced to 0 for infected mice that received 50 and 100 mg/kg MM3122 prophylactically (*P <* 0.01), and 0.5 for mice that received 50 and 100 mg/kg MM3122 therapeutically. Mice that received MM3122 prior to infection also had reduced virus titers (5000-10,000-fold) compared to vehicle treated and infected animals (**Fig 2B**). However, mice that received MM3122 24 h after infection had similar amounts of virus in the lungs compared to the vehicle treated animals. Finally, we performed pathological analysis of the lungs of MM3122 and control treated mice. Global pneumonia, used to assess the % of lung affected, was between 3 (>50%) and 4 (>80%) for the SARS-CoV-2 infected and vehicle treated mice. This score was reduced to 1.2 (*P <* 0.05) and 1.6 in the mice that received 50 mg/kg and 100 mg/kg of MM3122 prior to virus infection. No change in the global pneumonia score was observed for the mice that received 50 (score = 3) and 100 (score = 3.2) mg/kg of MM3122 after virus infection. Similarly, the bronchointerstitial pneumonia score was reduced in the animals that received MM3122 prior to infection (1.2 and 1.5 for 50 mg/kg (*P <* 0.05) and 100 mg/kg respectively) but not after SARS-CoV-2 infection. Finally, vasculitis and endotheliitis replicated control baseline levels (average score = 0.3) in the prophylactically treated group (score = 0.4 and 0.3 for 50 mg/kg and 100 mg/kg respectively). A reduction in score was also observed for the mice that received MM3122 after infection (score = 1.2 and 1.8 for 50 mg/kg and 100 mg/kg respectively), but this was not statistically significant compared to the infected and vehicle control treated animals (score = 2.8).

**Figure 2:**
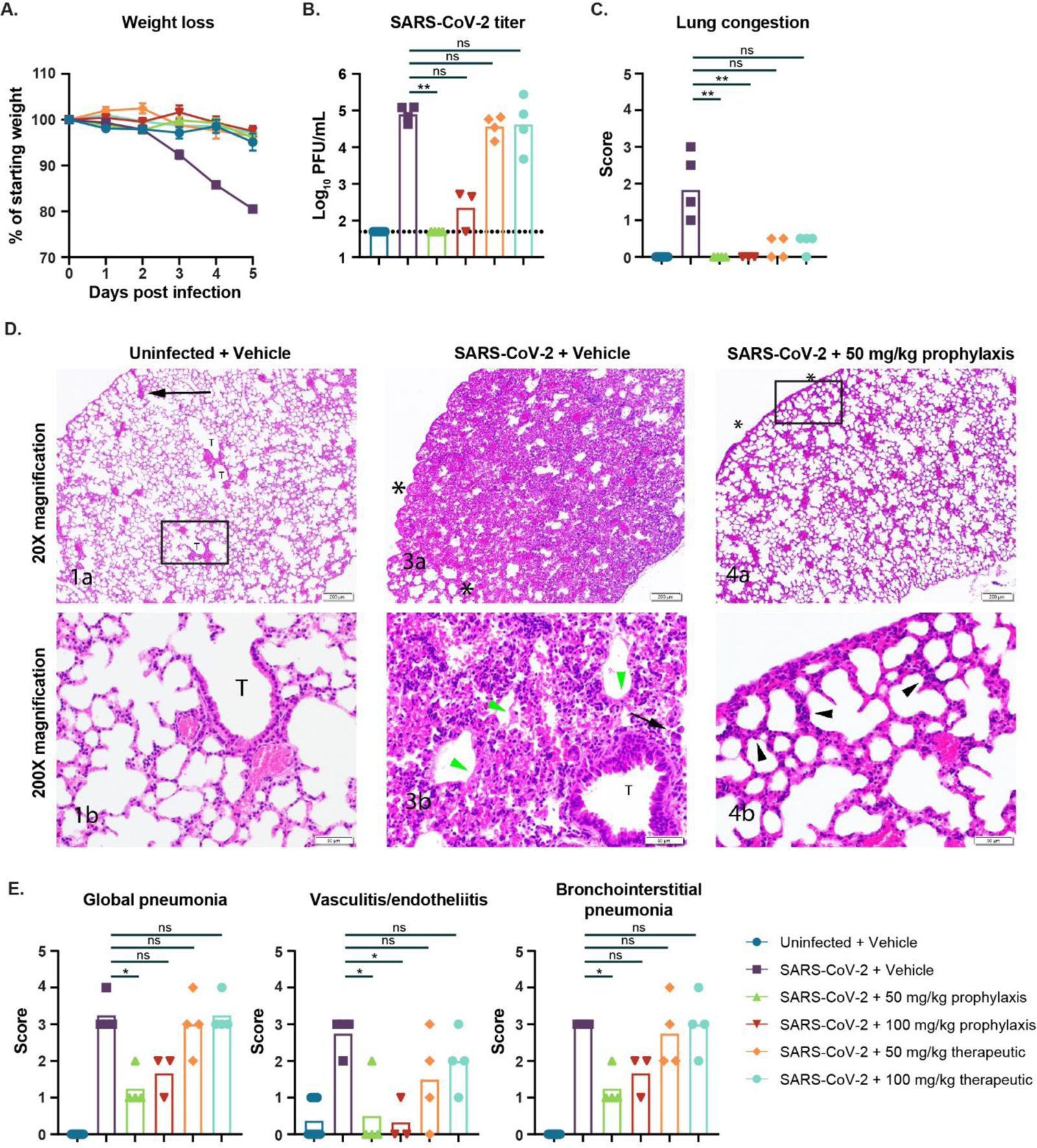
MM3122 protects mice against SARS-CoV-2 disease in mice. Eleven-to twelve-month-old female mice received MM3122 30 minutes before and 24 hours after intranasal inoculation with 1,000 PFU of MA10. (**A**) Weight loss was measured daily for five days. (**B**) Infectious virus titer in left lung lobe 5 days after infection. (**C**) Lung congestion score 5 days after infection. (**D**) Representative images and magnifications of H&E sections of lungs from uninfected and vehicle control treated mice (left panel), SARS-CoV-2 infected, and vehicle control treated mice (middle panels), and MM3122 treated and SARS-CoV-2 infected mice (right panels). (**E**) Global pneumonia, vasculitis and endotheliitis, and bronchointerstitial pathology scores of these same animals. All data was analyzed by a non-parametric one-way ANOVA (Kruskal-Wallis) with multiple comparisons corrects against the SARS-CoV-2 + vehicle group. The dotted line is the limit of detection. Each data point is an individual mouse, and the data are from a single experiment with 3-4 mice per group. ** = *P* < 0.01, ns = not significant.

### MM3122 reduces inflammatory cytokine and chemokine production after SARS-CoV-2 infection

Weight loss and severe disease after SARS-CoV-2 infection is associated with exacerbated inflammatory responses resulting in lung congestion and immunopathology. To assess inflammation, cytokine and chemokine concentrations were quantified in lung tissue homogenates using a mouse cytokine 23-plex assay. Compared to uninfected and vehicle control animals, the levels of IL-6, KC, G-CSF, CCL2, CCL3, CCL4, IL1 alpha, IL-12p40 and CCL5 were increased three-fold or more five days after infection with the MA10 strain of SARS-CoV-2 (**Fig 3**). Prophylactic and therapeutic treatment with MM3122 significantly (*P <* 0.001) reduced the amount of IL-6, KC, G-CSF, CCL2 and CCL3 in the lungs of these mice. Smaller reductions in cytokine and chemokine productions were observed for CCL4, IL-1 alpha, IL-12p40, and CCL5. Combined, these data show that the TMPRSS2 inhibitor MM3122 reduces virus titers and inflammation after SARS-CoV-2 infection.

**Figure 3:**
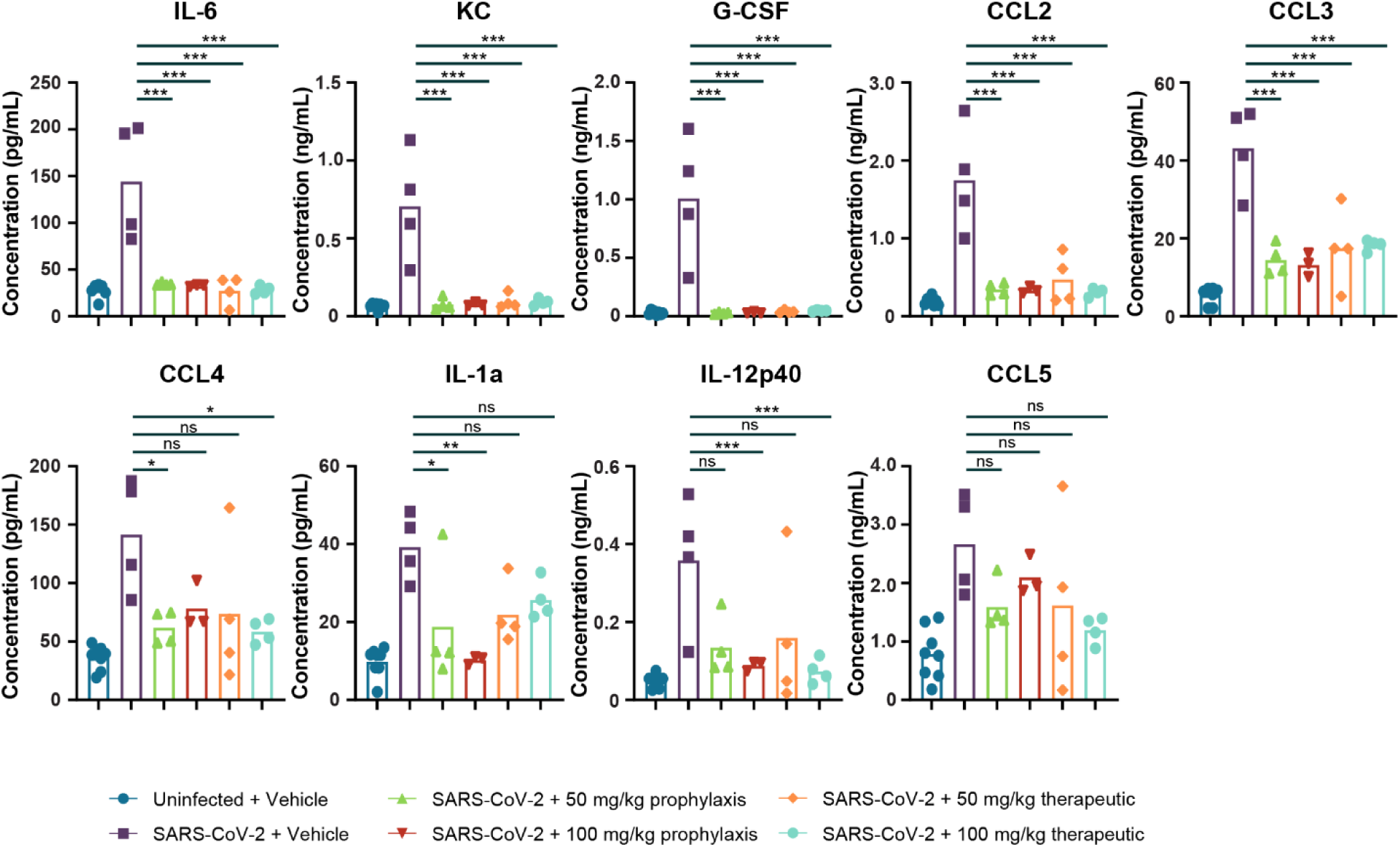
Prophylactic and therapeutic administration of MM3122 reduces inflammation in mice. Eleven-to twelve-month-old female mice received MM3122 30 minutes before and 24 hours after intranasal inoculation with 1,000 PFU of SARS-CoV-2 MA10. Cytokine and chemokine concentrations in left lung lobe homogenates collected from these same animals. Data was analyzed by a parametric ordinary one-way ANOVA with multiple comparisons corrects against the SARS-CoV-2 + vehicle group. Each data point is an individual mouse, and the data are from a single experiment with 3-4 mice per group. * = *P* < 0.05, ** = *P* < 0.01, *** = *P* < 0.001, ns = not significant.

## DISCUSSION

The COVID-19 pandemic began 4 years ago and is still a major economic catastrophe and medical problem worldwide costing over $14 trillion in economic losses in the US^45^ and resulting in excess mortality rates estimated at exceed 24 million people worldwide through early 2023^46^. While highly efficacious vaccines have saved millions of lives globally^47^, they are not 100% effective, even less so for variants, and a large percentage of the population refuses to be vaccinated so there is a dire need for new drugs to prevent and treat this life-threatening disease. There are some small molecule FDA-approved drugs to treat COVID-19^1^ including the viral polymerase inhibitors remdesivir and molnupiravir, as well as SARS-CoV-2 Mpro protease inhibitor nirmatrelvir, sold under the brand name Paxlovid. With the exception immunomodulatory treatments for COVID-19 such as the JAK1/JAK2 inhibitor baricitinib^48–50^, which target the symptoms of infection and not viral pathogenesis directly, there are no approved antiviral drugs, which target the host and confer protection against multiple different viruses from diverse viral families. Concomitantly, others, and we have reported on the first potential drugs, targeting the TMPRSS2 and matriptase, to treat SARS-CoV-2 and COVID-19^39, 42^. TMPRSS2, and other host cell transmembrane proteases^51^ including matriptase are essential for viral entry of many respiratory RNA viruses into the lung tissue. In this communication, we have demonstrated that one lead candidate drug that we have developed, MM3122, exhibited significant protective effects against weight loss, lung congestion (gross lung discoloration), and inflammation administered prophylactically at both low and high doses, in aged mice infected with mouse-adapted SARS-CoV-2. These effects were less pronounced in the therapeutic groups. Interestingly, these protections did not fully correspond with protection from viral replication, as titers were lower or absent in the prophylactic group but were not significantly different from the infected vehicle control in the therapeutic group. These protective effects of COVID-19 in the lung can potentially be explained by inhibition of the proteolytic activation of other substrates by TMPRSS2, matriptase, and hepsin or the other proteases, which MM3122 targets such as tryptase (**Table 1**). In summary, these promising results suggest MM3122 is a potential clinical candidate for the treatment and prevention of diseases including COVID-19, caused from infection by SARS-CoV-2 and other coronaviruses.

### Limitations of the study

We note several limitations of our study. (a) The *in vivo* efficacy of MM3122 was not tested against more recent variants of SARS-CoV-2 in mice or in Syrian hamsters. For the latter, we will need to perform PK studies, which is beyond the scope of this study. Also, the more recent variants of SARS-CoV-2 are attenuated compared to MA10 strain of SARS-CoV-2, creating challenges in observing effects on weight loss and disease of MM3122. Also, given that MM3122 has similar activity against EG.5.1 *in vitro*, we do not expect differences *in vivo*. (b) We did not evaluate the efficacy of MM3122 in male mice. Males are considered more susceptible to SARS-CoV-2 infection and therefore we expect more disease in the untreated animals with continued amelioration of disease in the MM3122 treated animals. (c) To increase efficacy, we are currently working on developing improved compounds with longer half-life and higher compound exposure compared to MM3122.^39^ One potential way to achieve increased efficacy with MM3122 would be to develop formulations for inhalation or nebulization as alternative administration routes, which would deliver compound directly to the respiratory tract at the primary site of viral entry and the infection. In summary, these studies demonstrate that MM3122 is effective in inhibiting SARS-CoV-2 replication *in vitro* and that administration of MM3122 *in vivo* reduces COVID-19 disease in mice.

## MATERIALS AND METHODS

### Cells and Viruses

Vero cells expressing human angiotensin converting enzyme 2 (ACE2) and transmembrane protease serine 2 (TMPRSS2) (Vero-hACE2-hTMPRSS2^52, 53^, gift from Adrian Creanga and Barney Graham, NIH) were cultured at 37°C in Dulbecco’s Modified Eagle medium (DMEM) supplemented with 10% fetal bovine serum (FBS), 10 mM HEPES (pH 7.3), 100 U/mL of Penicillin, 100 µg/mL of Streptomycin, and 10 µg/mL of puromycin. Vero cells expressing TMPRSS2 (Vero-hTMPRSS2)^53^ were cultured at 37°C in DMEM supplemented with 10% fetal bovine serum (FBS), 10 mM HEPES (pH 7.3), 100 U/mL of Penicillin, 100µg/mL of Streptomycin, and 5 µg/mL of blasticidin. Calu-3 cells were cultured in DMEM media supplemented with 1.0 mM sodium pyruvate, non-essential amino-acids (NEAA), 100 U/mL of penicillin, 100 µg/mL streptomycin, 2.0 mM L-glutamine, 10 mM HEPES, and 10% Fetal Bovine Serum (FBS).

The Lineage A variant of SARS-CoV-2 (WA1/2020), or the XBB.1.5 and E.G.5.1 (from Mehul Suthar) variants of SARS-CoV-2 were propagated on Vero-hTMPRSS2 cells. The virus stocks were subjected to next-generation sequencing, and the S protein sequences were identical to the original isolates. The infectious virus titer was determined by plaque and focus-forming assay on Vero-hACE2-hTMPRSS2 or Vero-hTMPRSS2 cells.

Baric laboratory-generated stock of SARS-CoV-2 MA10, a mouse-adapted virulent mutant generated from a recombinantly derived synthesized sequence of the Washington strain that causes severe acute and chronic disease in mice^54, 55^. Virus was maintained at low passage (P2-P3) to prevent the accumulation of additional potentially confounding mutations.

### Drug Preparation and Administration

MM3122^39^ was freshly prepared at 8 mg/mL in 5% DMSO in PBS. MM3122 was administered intraperitoneal (IP) at 50 and 100 mg/kg bodyweight in 100µL volumes to mice starting at 30 minutes before infection (prophylactic treatment) or 24 hours after infection (day 1, therapeutic treatment). Subsequent doses were administered at approximately the same times each day post-infection.

### SARS-CoV-2 challenge studies

All studies with mice were conducted under the University of North Carolina IACUC approval (20-114). Aged (11-to 12-month-old) female BALB/c mice obtained from Envigo (retired breeders) were acclimated for 7 days in the Biosafety laboratory level 3 prior to any experimentation. Food and water were provided ad libitum, and the animal room maintained a 12-hour light/dark cycle. Prior to inoculation with SARS-CoV-2, animals were anesthetized intraperitoneal with a combination of 50 mg/kg Ketamine and 15 mg/kg Xylazine in 50 µL, and infected intranasally with 1,000 PFU of sequence-and titer-verified SARS-CoV-2 MA10 in 50 µL PBS. Mice were monitored daily for weight loss and disease. At 5 days post-infection, mice were euthanized following sedation by isoflurane and thoracotomy, and lungs were collected for assessments of virus titer, inflammatory cytokines and chemokine levels, histological analysis, and lung congestion score. Lung congestion score was measured using an independently defined scale of 0-4 (0: no congestion; 1: one lobe involved; 2: two lobes involved; 3: three lobes involved; 4: all four lobes involved; all scores have 0.5-point intervals). Lung sections used in all experiments for various assessments were as follows: lower right lobe, histology; upper right lobe, RNA; left lobe and central lobe, virus titer. Prior to virus titration and cytokine analysis, the lung lobes were homogenized with glass beads in 1.0 mL PBS, clarified by centrifugation and stored at -80°C. Infectious virus titers were quantified by plaque assay on Vero E6 cells and calculated at PFU/mL of homogenized tissue.

### Cytokine Analysis

Homogenized lung samples were subjected to cytokine and chemokine analysis using the Bio-Rad Mouse Cytokine 23-plex assay (Cat # M60009RDPD) per the manufacturer’s protocol. Cytokine assays were performed in non-inactivated samples at BSL3. The data were analyzed on a Luminex MAGPIX machine, and cytokine concentrations in the lung homogenates were extrapolated using the provided standards.

### Histological analysis

Lung tissues from the lower right lobe were fixed for a minimum of 7 days in 10% formalin, paraffin embedded, sectioned, and stained with hematoxylin and eosin (H&E). H&E sections were submitted for graded blindly for vasculitis/endotheliitis, Bronchointerstitial pneumonia, and global pneumonia severity score by a board-certified veterinary pathologist. Details on the histology scoring system are provided in the Supplementary Material.

### MM3122 in vitro inhibition assays

Calu-3 cells (5 x 10^5^ cells/well) were seeded in 24-well culture plates in infection medium (DMEM + 1.0 mM Sodium pyruvate, NEAA, 100 U/mL of penicillin, 100 µg/mL streptomycin, 2.0 mM L-glutamine, 10 mM HEPES, and 2% FBS) and incubated overnight at 37°C and 5% CO_2_. After 24 h, media was removed and fresh 250 µL media was added to each well containing MM3122 or Remdesivir starting at 60 µM concentration and diluted 3-fold to 20, 6, 2, 0.6 and 0.2 µM. Media alone, and DMSO were included as negative controls. Next, the cells were transferred to the BSL3 laboratory and 250 µL of media containing 4,000 PFU of SARS-CoV-2 was added for 1 h at 37°C and 5% CO_2_. Note, that the final concentration of MM3122 and Remdesivir is 30, 10, 3, 1, 0.3, and 0.1 µM. After 1 h, the virus inoculum was removed, the cells were washed twice with infection media and fresh infection media containing MM3122, Remdesivir or DMSO was added to each well. At 48 h post-infection, culture supernatant is collected and used to quantify virus titers by plaque assay as described below.

### Virus titration assays

Plaque assays were performed on Vero-hACE2-hTRMPSS2 cells in 24-well plates. Lung tissue homogenates or nasal washes were diluted serially by 10-fold, starting at 1:10, in cell infection medium (DMEM + 100 U/mL of penicillin, 100 µg/mL streptomycin, and 2% FBS). Two hundred and fifty microliters of the diluted virus were added to a single well per dilution per sample. After 1 h at 37°C, the inoculum was aspirated, the cells were washed with PBS, and a 1% methylcellulose overlay in MEM supplemented with 2% FBS was added. Seventy-two to ninety-six hours after virus inoculation, dependent on the virus strain, the cells were fixed with 4% formalin and the monolayer was stained with crystal violet (0.5% w/v in 25% methanol in water) for 30 min at 20°C. The number of plaques were counted and used to calculate the plaque forming units/mL (PFU/mL).

## ACKNOWLEDGEMENTS

This study was supported by the NIH (NIAID Center of Excellence for Influenza Research and Response (CEIRR)) contract 75N93021C00016 and R01 AI169022 to A.C.M.B and by Washington University School of Medicine, Siteman Cancer Center grant SCC P30CA091842 and Barnes Jewish Hospital Foundation award BJHF 4984 to J.W.J. We also acknowledge preclinical services provided by the National Institute of Allergy and Infectious Diseases, National Institutes of Health, Department of Health and Human Services, under contract HHSN272201700036I/75N93020F00001 to the University of North Carolina-Chapel Hill.

## AUTHOR CONTRIBUTIONS

T.L.B. performed the *in vitro* inhibition studies on Calu-3 cells. E.J.F., S.R.L., K.G., R.S.B, R.L.G. performed all the *in vivo* mouse studies with MM3122 and SARS-CoV-2. B.V.T performed the histology experiments. M.M. synthesized the MM3122 compound. T.L.B., A.C.M.B., R.L.G., R.S.B., J.W.J. analyzed the data. A.C.M.B. performed the statistical analysis. A.C.M.B. had unrestricted access to all the data. A.C.M.B, J.W.J. provided key reagents, supervised experiments, and acquired funding. A.C.M.B. and J.W.J. wrote the manuscript and all authors reviewed and edited the final version. All authors agreed to submit the manuscript, read, and approved the final draft, and take full responsibility of its content.

## SUPPLEMENTARY INFORMATION

Scoring criteria for Global pneumonia severity.

| Grade | Criteria |
| --- | --- |
| Normal = 0 | The lung is considered to be within normal limits, under the conditions of the study and considering the age, sex, and strain of the animal |
| Minimal = 1 | ≤ 25% of lung affected |
| Mild = 2 | 26-50% of lung affected |
| Moderate = 3 | 51-79% of lung affected |
| Marked = 4 | ≥ 80% of lung affected |

Scoring criteria for Vasculitis / Endotheliitis.

| Grade | Criteria |
| --- | --- |
| Normal = 0 | No significant microscopic changes present above background lesions if any. |
| Minimal = 1 | Multifocal, scattered, individual infiltrates of inflammatory cells are present along the surface, below the endothelium, or within the tunica media or tunica adventitia of rare vessels. |
| Mild = 2 | Infiltrates of small numbers of individual to occasional focal aggregates of inflammatory cells are present along the surface, below the endothelium, or within the tunica media or tunica adventitia of multiple vessels. There is rare leukocytoclastic vasculitis. |
| Moderate = 3 | Infiltrates of variable numbers of inflammatory cells are present along the surface, below the endothelium, or within the tunica media or tunica adventitia of the majority of vessels. Focal aggregates are common within the endothelium narrowing the lumen and there is occasional leukocytoclastic vasculitis. |
| Marked = 4 | Infiltrates of focally large numbers of inflammatory cells are present along the surface, below the endothelium, or within the tunica media or tunica adventitia of almost all vessels. Large aggregates are common within the endothelium narrowing the lumen and there is common leukocytoclastic vasculitis. |

Scoring criteria for Bronchointerstitial pneumonia.

| **Grade** | **Criteria** |
| --- | --- |
| Normal = 0 | No significant microscopic changes present above background lesions, if any. |
| Minimal = 1 | Affected alveolar septa are visibly but minimally thickened and there are only small amounts of hyaline membrane formation, edema, fibrin deposition, or hemorrhage within bronchioles and alveolar airways, with little loss of total airway space. |
| Mild = 2 | Affected alveolar septa are mildly thickened and hyaline membrane formation, edema, fibrin deposition, or hemorrhage are commonly observed within alveolar spaces and bronchioles. Focal areas of atelectasis and consolidation may be present. |
| Moderate = 3 | Infiltrates of inflammatory cells are present within the septa and spaces of most alveoli. Affected alveolar septa are moderately thickened, and hyaline membrane formation, edema, fibrin deposition, or hemorrhage are frequently observed within alveolar spaces or bronchioles along with prominent loss of airspace. Multifocal and regional areas of atelectasis and consolidation are present. |
| Marked = 4 | Infiltrates of large numbers of inflammatory cells are present within the septa and spaces of most alveoli. Affected alveolar septa are markedly thickened, and hyaline membrane formation, edema, fibrin deposition, or hemorrhage within alveolar airways and bronchioles are frequently observed. Large areas of atelectasis and consolidation are present. |

